# Targeted Next-Generation Sequencing Identifies Pathogenic Variants in Kidney Disease-Related Genes in Patients with Diabetic Kidney Disease

**DOI:** 10.1101/2020.05.18.102822

**Authors:** Jose Lazaro-Guevara, Julio Fierro Morales, A. Hunter Wright, River Gunville, Scott G. Frodsham, Melissa H. Pezzolesi, Courtney A. Zaffino, Laith Al-Rabadi, Nirupama Ramkumar, Marcus G. Pezzolesi

## Abstract

Diabetes is the most common cause of chronic kidney disease (CKD). For patients with diabetes and CKD, the underlying cause of their kidney disease is often assumed to be a consequence of their diabetes. Without histopathological confirmation, however, the underlying cause of their kidney disease is unclear. Recent studies have shown that next-generation sequencing (NGS) provides a promising avenue toward uncovering and establishing precise genetic diagnoses in various forms of kidney disease. Here, we set out to investigate the genetic basis of disease in non-diabetic kidney disease (NDKD) and diabetic kidney disease (DKD) patients by performing targeted NGS using a custom panel comprised of 345 kidney disease-related genes. Our analysis identified rare diagnostic variants that were consistent with the clinical diagnosis of 19% of the NDKD patients included in this study. Similarly, 22% of DKD patients were found to carry rare pathogenic/likely pathogenic variants in kidney disease-related genes included on our panel. Genetic variants suggestive of NDKD were detected in 3% of the diabetic patients included in this study. Our findings suggest that rare variants in kidney disease-related genes in the context of diabetic pathophysiology may play a role in the pathogenesis of kidney disease in patients with diabetes.

**Key Messages:**

1. What is already known about this subject?

- For patients with diabetes and chronic kidney disease, the underlying cause of their kidney disease is often assumed to be a consequence of their diabetes; without histopathological confirmation, however, the underlying cause of their kidney disease is unclear.
- Next-generation sequencing (NGS) provides a promising avenue toward uncovering and establishing precise genetic diagnoses in various forms of kidney disease.
2. What are the new findings?

- Using targeted NGS and a custom panel comprised of 345 kidney disease-related genes, we found that 22% of diabetic kidney disease patients were found to carry rare pathogenic/likely pathogenic variants in kidney disease-related genes included on our panel.
- Genetic variants suggestive of non-diabetic kidney disease were detected in 3% of the diabetic patients included in this study.
3. How might these results change the focus of research or clinical practice?

- Our findings suggest that rare variants in kidney disease-related genes in the context of diabetic pathophysiology may play a role in the pathogenesis of kidney disease in patients with diabetes.
- Importantly, improved understanding of the underlying disease process in diabetic kidney disease could have major implications in terms of patient care and monitoring as well as for research studies in this field.

## Introduction

Diabetes is the most common cause of chronic kidney disease (CKD) (1). Continued increase in the world-wide prevalence of diabetes has led to an increase in the global prevalence of diabetic kidney disease (DKD), also known as diabetic nephropathy (2). For patients with diabetes and CKD, the underlying cause of their kidney disease is often assumed to be a consequence of their diabetes. However, without histopathological evidence, it’s unclear whether such patients have true DKD, non-diabetic kidney disease (NDKD), or concomitant DKD and NDKD.

The only way to truly determine whether CKD in patients with diabetes is a direct consequence of diabetes is to perform renal biopsies. Unfortunately, renal biopsies are not clinically indicated in the diagnosis of DKD. Interestingly, recent investigations of renal biopsies from patients with type 1 diabetes (T1D) or type 2 diabetes (T2D) and CKD have shown that as many as 30-83% of patients diagnosed with DKD actually had kidney disease attributed to non-diabetic causes (3–5). Familial focal segmental glomerulosclerosis (FSGS), followed by hypertensive nephropathy, acute tubular necrosis, and IgA nephropathy were the most common diagnoses observed in diabetic patients found to have NDKD (6). Among T1D patients, the prevalence of NDKD is estimated to be 2-3%, while more than 60% of T2D patients are likely to have NDKD with or without DKD (6). Importantly, accurately identifying a patient’s primary cause of CKD is a crucial component of its proper classification, prognosis, and management.

Recent studies conducted primarily in patients with NDKD have shown that next-generation sequencing (NGS) provides a promising avenue toward uncovering and establishing precise genetic diagnoses in various forms of kidney disease (7–12). The diagnostic yield from these studies range from approximately 10-40% and are highest among patients with congenital or cystic renal disease. While these studies all show the utility of NGS in providing a molecular diagnosis for patients with heritable forms of kidney disease, whether this technology could also aid in the genetic diagnosis of patients with DKD is unclear. Importantly, improved understanding of the underlying disease process in DKD could have major implications in terms of patient care and monitoring as well as for research studies in this field. To date, no study has examined the distribution of rare variants in known kidney disease-related genes in patients with DKD. To address this, we performed targeted NGS using a custom gene panel comprised of 345 kidney disease-related genes in 222 patients with CKD, including 98 NDKD and 124 DKD patients.

## Methods

### Study Cohorts

A total of 222 patients with CKD, including 98 non-diabetic patients (NDKD) and 124 diabetic (DKD) patients, were recruited to the Utah Kidney Study from participating nephrology dialysis centers in the University of Utah Health Center (UUHSC) network between January 1, 2017 and December 31, 2018. All patients provided detailed medical and family history information through patient questionnaires. All diabetic patients included in this study had a diagnosis of diabetes that predated their CKD diagnosis and reported diabetes as the primary cause of their CKD. Additional medical information, including CKD-related diagnoses, diabetes status, and measures of kidney function (i.e., serum creatinine and eGFR measurements), were obtained through the UUHSC’s electronical medical records (EMRs). Blood and urine samples were obtained at time of recruitment. Serum creatinine measurements, determined at ARUP Laboratories (Salt Lake City, UT) by standard enzymatic methods, and the Chronic Kidney Disease Epidemiology Collaboration (CKD-EPI) (13) were used to estimated GFR at the time of recruitment. Genomic DNA was isolated from blood samples using a standard phenol:chloroform DNA extraction protocol. Written and informed consent was obtained from all participants. This study was approved by the University of Utah Institutional Review Board (IRB_00098113 and IRB_00109765).

### Kidney Disease-related Gene Panel and Targeted Next-Generation Sequencing

A custom gene panel was designed for simultaneous interrogation of genetic variation across 345 kidney disease-related genes (**Supplemental Table 1**). Kidney disease-related genes were selected using the Human Genome Database (www.hgmd.com), Online Mendelian Inheritance in Man (OMIM; www.omim.org) and a comprehensive literature review. Genomic positions of the coding sequence of all 345 genes were obtained from the Consensus Coding Sequence database (www.ncbi.nlm.nih.gov/CCDS.CcdsBrowse.cgi), Release 20). All exons and exon-intron boundaries of these genes were captured using a custom capture assay (Agilent Technologies, Santa Clara, CA). The targeted sub-genome spanned a total of 972,494 basepairs. Samples were fragmented using a Covaris S2 ultrasonicator (Covaris, Woburn, MA) and prepared for sequencing on an Illumina HiSeq2500 using a custom DNA library preparation protocol based on the method described by Rohland et al. (14).

Following sequencing, the resulting reads were aligned to the human reference genome (hg19) with Sentieon BWA (Sentieon, Mountain View, CA). Evaluation of the aligned reads and variant calling was performed using the Sentieon DNAseq pipeline. Six samples were excluded from variant calling and downstream analysis due to insufficient read depth. The average read depth across the 216 remaining samples was 309X with 98.1% of bases above 15X coverage. We then inferred family relationships among these patients using ~9,000 informative SNPs from the targeted sequencing data and the Kinship-based INference for Gwas (KING) software (15). We identified 10 first-degree relative pairs (kinship coefficient > 0.125). Following removal of one member from each of these relative pairs, the final NDKD and DKD cohorts included 97 and 109 patients, respectively. Clinical characteristics for these patients are presented in **Table 1**.

**Table 1:** Clinical Characteristics of NDKD and DKD Cohorts.

| Characteristic | NDKD | DKD |
| --- | --- | --- |
| N | 97 (47.1) | 109 (52.9) |
| Men | 38 (39.2) | 67 (61.5) |
| White | 73 (75.3) | 72 (75.0) |
| Age (years) | 47.9 (36.0, 60.0) | 62.0 (53.0, 68.0) |
| BMI | 28.0 (22.8, 34.1) | 31.4 (25.7, 36.0) |
| CKD Clinical Diagnosis: |  |  |
| Congenital Renal Disease | 5 (5.2) | --- |
| Cystic Renal Disease | 20 (20.6) | --- |
| Glomerulopathy | 55 (56.7) | --- |
| Tubulointerstitial Disease | 4 (4.1) | --- |
| Diabetic Nephropathy | --- | 109 (100.0) |
| Other | 13 (13.4) | --- |
| Type 2 Diabetes | 17 (17.5)* | 88 (80.7) |
| HbA1c (%) | 7.7 (6.5, 8.4) | 7.0 (6.4, 8.2) |
| Treatment with Insulin | 0 (0.0) | 83 (76.1) |
| Duration of Diabetes Prior to<br>Kidney Disease Diagnosis (years) | --- | 17.0 (10.0, 25.5) |
| eGFR at Exam (mL/min/1.73 m<sup>2</sup>) | 27.0 (10.0, 53.0) | 10.0 (10.0, 37.0) |
| CKD Stage at Exam: |  |  |
| CKD 1 (eGFR ≥ 90 mL/min/1.73 m<sup>2</sup>) | 9 (9.3) | 1 (0.9) |
| CKD 2 (eGFR 60-89 mL/min/1.73 m<sup>2</sup>) | 14 (14.4) | 7 (6.4) |
| CKD 3 (eGFR 30-59 mL/min/1.73 m<sup>2</sup>) | 22 (22.7) | 25 (22.9) |
| CKD 4 (eGFR 15-29 mL/min/1.73 m<sup>2</sup>) | 11 (11.3) | 3 (2.8) |
| CKD 5 (eGFR < 15 mL/min/1.73 m<sup>2</sup>) | 41 (42.3) | 73 (67.0) |
| First Degree Relative with Kidney Disease | 37 (38.1) | 37 (33.9) |

Data are presented as *n* (%) or median (1st, 3rd quartile). HbA1c, hemoglobin A1c; CKD, chronic kidney disease; BMI, body mass index; eGFR, estimated glomerular filtration rate.
\*17 NDKD patients were diagnosed with type 2 diabetes after developing CKD.

### Variant Analysis

The resulting variant call format (VCF) file was annotated using ANNOVAR (16) and filtered based on variant function (nonsynonymous, stop gain/loss, splicing, frameshift insertion/deletion, and in-frame insertion/deletion variants) and minor allele frequencies (MAF; < 0.1% global population MAF) in the Genome Aggregation Database (gnomAD; gnomad.broadinstitute.org/) (17), a population-level database of genomic variant frequencies derived from large-scale exome and genome sequencing data. The clinical significance of these candidate rare, functional variants was interpreted using InterVar (18) following the 2015 American College of Medical Genetics and Genomics and the Association for Molecular Pathology (ACMG-AMP) standards and guidelines (19). Variants classified as ‘pathogenic’ or ‘likely pathogenic’ according to these guidelines and that followed the inheritance pattern (dominant or recessive) associated with disease were considered to be diagnostic variants causal of the patient’s nephropathy. Although phasing information was not available, we considered individuals with at least 2 rare (MAF < 0.1%) heterozygous genotypes in a single gene to be putative compound heterozygous carriers under a recessive model. In addition, multiple computational prediction algorithms, including SIFT (20), Polyphen2 (Polyphen2_HDIV and Polyphen2_HVAR) (21), MutationTaster (22), M-CAP (23) and LRT (24), were utilized to aid in the interpretation of nonsynonymous variants classified as ‘variants of unknown significance’ (VUSs) by the ACMG-AMP guidelines. In line with these guidelines, supportive evidence of pathogenicity was considered for VUSs predicted to be damaging by at least 5 of these 6 prediction methods.

Additionally, we performed copy-number variant (CNV) analysis using Atlas-CNV, a method for detecting and prioritizing high-confidence CNVs in targeted NGS data (25). As recommended by the developers of this program, CNVs with a log_2_ score −0.6 (losses) and 0.4 (gains), corresponding to confidence scores (C-scores) of −7.49 and 5.01, respectively, were called. All prioritized CNVs were visualized using the Integrative Genomics Viewer software program (26) and inspected manually.

### Statistical Analysis

Continuous data are presented as the median (1^st^, 3^rd^ quartile). Dichotomous data are shown as n (percentage). Differences between the NDKD and DKD groups were assessed using χ^2^ tests and Fisher’s Exact Test (or Test for Trend) for dichotomous variables and unpaired *t* tests for continuous data comparisons using SAS software version 9.4 (SAS Institute, Cary, NC). Gene-based rare variant association burden testing of identified variants using the SNP-set (Sequence) Kernel Association Test (SKAT) (27). Two-tailed *p*-values less than 0.05 were considered statistically significant.

## Results

### Diagnostic Variants in Kidney Disease-related Genes in NDKD Patients

Using our targeted sequencing panel, we identified 1,259 rare, functional variants (i.e., nonsynonymous, stop gain/loss, splicing, frameshift insertion/deletion, and in-frame insertion/deletion variants) in 345 kidney disease-related genes among 206 patients included in this study (**Supplemental Table 2** and **Supplemental Table 3**). Of these variants, 63 (5.0%) were classified as pathogenic or likely pathogenic using the ACMG-AMP guidelines; the remaining variants were classified as VUSs (n=1,024; 81.3%) or benign/likely benign variants (n=172; 13.7%).

**Table 2:**
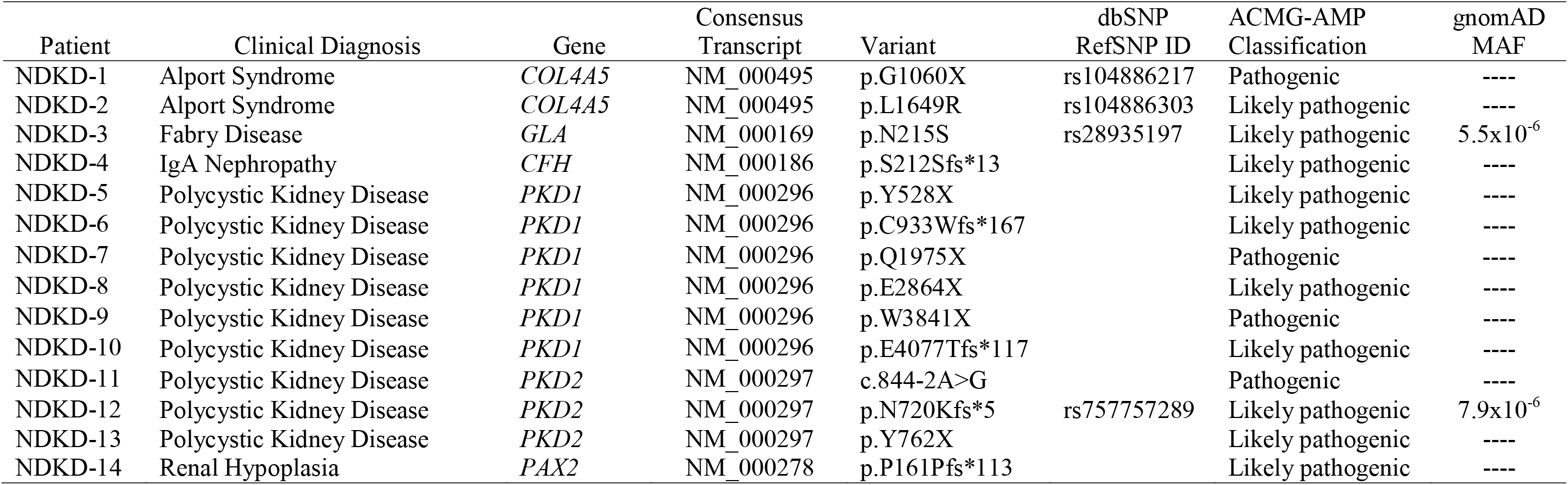
Summary of Diagnostic Variants Identified in NDKD Patients.

In total, pathogenic or likely pathogenic variants were identified in 35 of the 97 NDKD patients (36%) included in our study (**Supplemental Table 4**). We identified diagnostic variants (i.e., pathogenic or likely pathogenic variants consistent with the inheritance pattern associated with the patient’s clinical diagnosis) in 18 of the 97 NDKD patients (18.6%) included in our study (**Table 2** and **Table 3**). In total, 16 pathogenic or likely pathogenic single nucleotide and insertion/deletion variants, were detected in 7 genes, including 12 (75.0%) novel variants not present in gnomAD. *PKD1* and *PKD2* were the genes with the highest burden of candidate diagnostic variants (**Figure 1**).

**Table 3:**
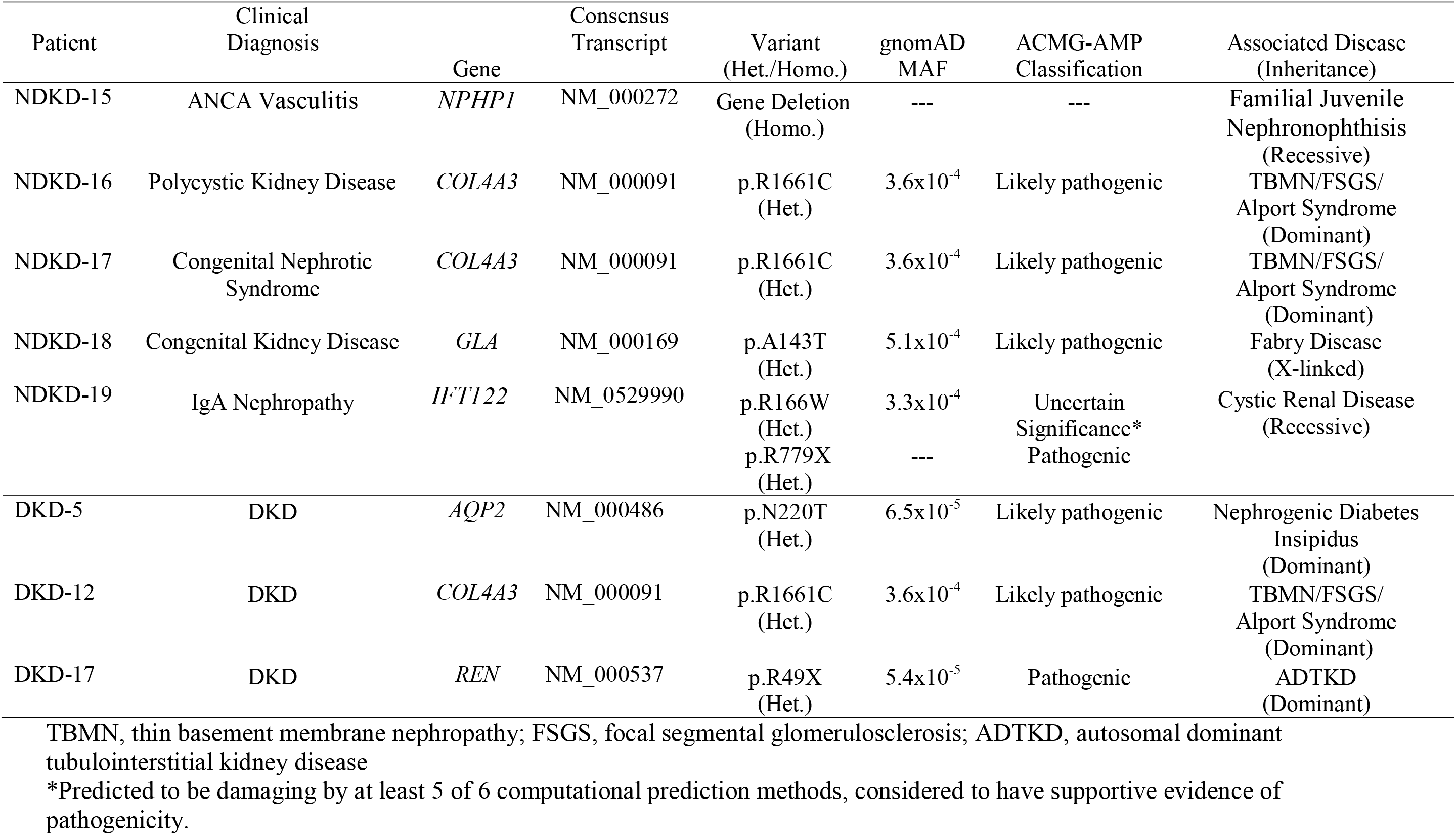
Re-classification of Clinical Diagnosis of NDKD and DKD Patients Based on Identified Diagnostic Variants.

**Figure 1.**
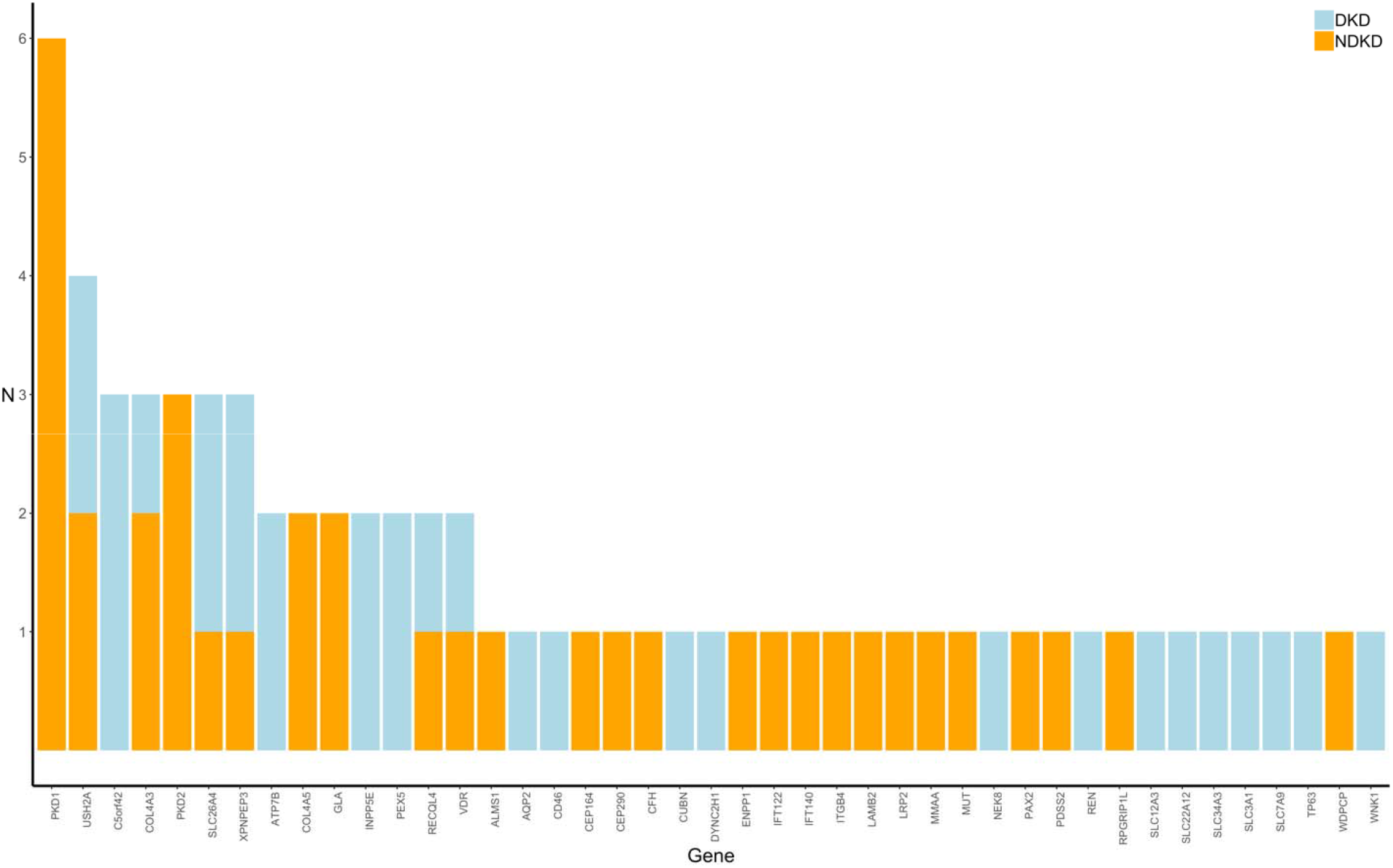
Distribution of Rare Pathogenic or Likely Pathogenic Variants in NDKD and DKD Patients. The distribution of rare (MAF < 0.1%) pathogenic or likely pathogenic variants are presented for NDKD (blue) and DKD (orange) patients.

Additionally, we detected a large homozygous deletion of *NPHP1* that has previously been shown to be pathogenic (28; 29) in one NDKD patient (**Supplemental Figure 1**). Other than a common deletion spanning the *CFHR3* and *CHFR1* genes (30), observed in 5 NDKD patients and 6 DKD patients, no additional structural variations, including inversions, translocations, or genomic imbalances (insertions and deletions), were observed.

The diagnostic yield was highest (9 of 20; 45.0%) in NDKD patients diagnosed with cystic renal disease. A total of 9 patients carried diagnostic variants in *PKD1* or *PKD2*, genes known to be causal of autosomal dominant polycystic kidney disease (ADPKD); 8 of the identified *PKD1/PKD2* variants were absent from population data. Five additional patients with cystic renal disease had novel VUSs in *PKD1* or *PKD2* that are predicted to be damaging using computational prediction tools (**Supplemental Table 5**). Although these variants do not meet strict ACMG-AMP criteria for classifying these as clinically significant, they do have supportive evidence of pathogenicity. Further examination of these variants, including segregation analysis and functional assays, together with this computational support, could increase the diagnostic yield of genetic screening in these cystic renal disease patients to as much as 70%.

### Re-Classification of Kidney Disease Diagnosis in NDKD Patients Based on a Genetic Diagnosis

Sequencing of kidney disease-related genes in 4 NDKD patients (4.1%) allowed us to identify variants that are consistent genetic diagnoses that differed from their reported clinical diagnoses (**Table 3**). Among these patients, we identified a large homozygous deletion of *NPHP1* in a patient diagnosed with antineutrophil cytoplasmic antibody-associated (ANCA) vasculitis following a renal biopsy. Interestingly, this well-characterized deletion is frequently seen in familial juvenile nephronophthisis (OMIM 256100), accounting for approximately 80% of patients with this disease (28; 29). Similarly, 1 patient diagnosed with PKD was found to have a diagnostic variant in *COL4A3* (p.R1661C). This same variant, although rare in the population (MAF = 3.6×10^-4^), was observed in 2 additional patients in our study, a patient with congenital nephrotic syndrome and a patient diagnosed with DKD, and has previously been observed in patients with FSGS (31) and autosomal recessive Alport syndrome (MIM 104200) (32). For each of these patients, their genetic diagnosis is more consistent with glomerular disease, including thin basement membrane nephropathy (TBMN), FSGS, or Alport syndrome. Similarly, we identified a rare variant in *GLA* (p.A143T) in a patient clinically diagnosed with congenital kidney disease; this variant has previously been reported to be pathogenic in patients with Fabry disease (MIM 301500) (33). Lastly, 1 NDKD patient diagnosed with IgA nephropathy was found to carry rare compound heterozygous variants in *IFT122* (p.R166W and p.R779X), which are known to contribute to ciliary dysfunction and cystic renal disease (34).

### Rare Variants in Kidney Disease-related Genes in Patients Diagnosed with DKD

Using the ACMG-AMP guidelines to classify variants in these same kidney disease-related genes, pathogenic or likely pathogenic variants were identified in 22.0% (24 of 109) of the DKD patients included in this study (**Table 4**). The proportion of variants was similar among both T1D and T2D patients (19% versus 23%, respectively; *p*-value > 0.05). Although not statistically significant (*p*-value > 0.05), nearly one-third of DKD patients (27.0%) found to carry a pathogenic or likely pathogenic variant reported a family history of kidney disease, compared to only 19.4% of DKD patients without a known family of kidney disease. In total, 29 pathogenic or likely pathogenic variants were identified in 23 kidney disease-related genes in these patients (**Figure 1**). *C5orf42* (also called *CPLANE1* and *JBTS17*) had the highest burden of pathogenic or likely pathogenic variants among these patients.

**Table 4:**
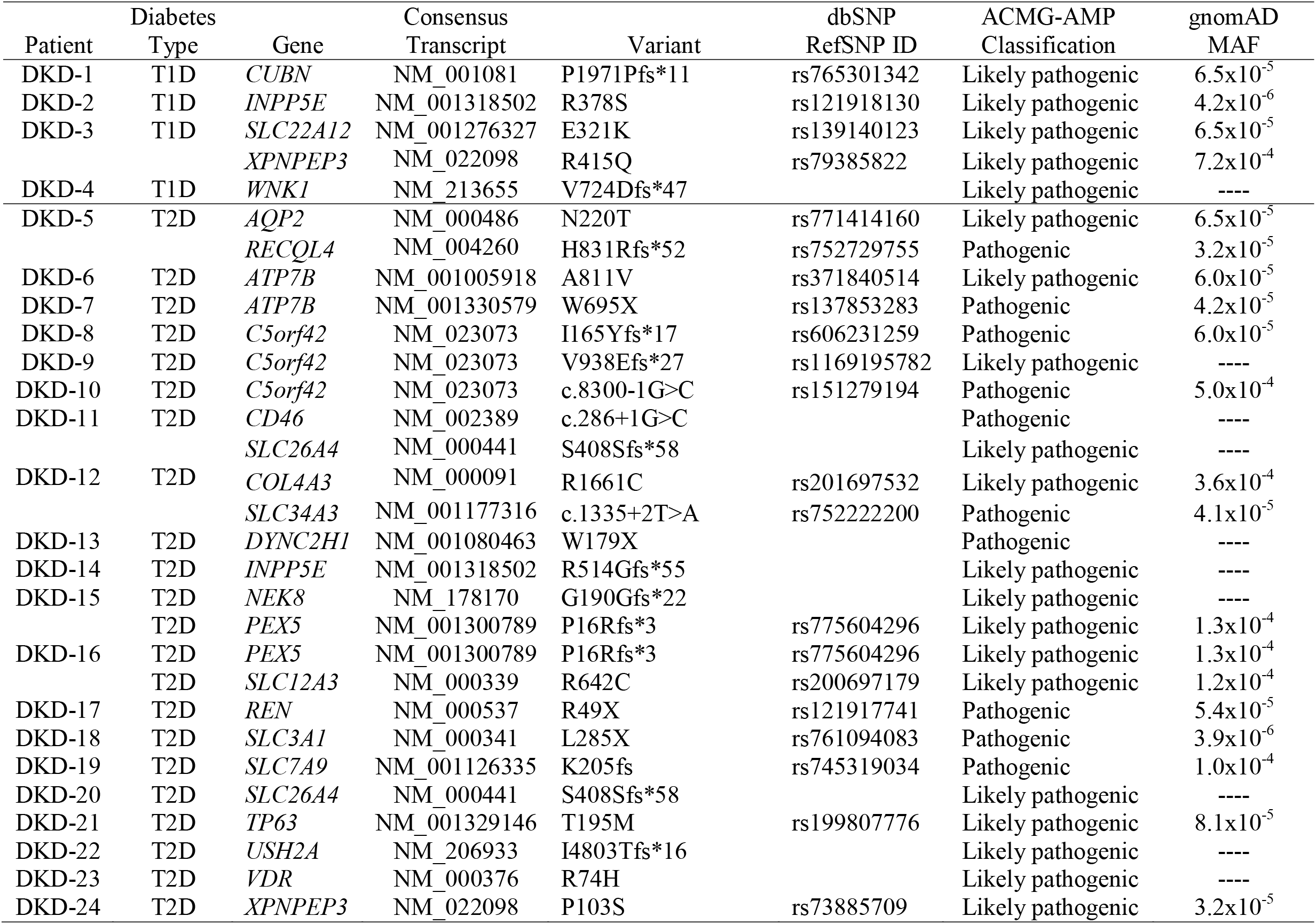
Summary of Pathogenic of Likely Pathogenic Variants in DKD Patients.

Interestingly, 9 DKD patients were found to harbor rare pathogenic or likely pathogenic variants in known ciliopathy genes, including *DYNC2H1, INPP5E, XPNPEP3, NEK8*, and *C5orf42*, and 6 of these variants are putative loss-of-function (pLOF) variants that affect splice sites or introduce premature stop codons (i.e., nonsense variants or frameshift insertion/deletions). Among these patients, 1 had biallelic variants in *C5orf42* (p.I165Yfs*17, a pathogenic variant, and p.S123F, a variant with supportive evidence of pathogenicity predicted to be damaging by at least 5 of 6 computational prediction methods), suggesting that this patient is likely a compound heterozygous carrier of damaging variants in this gene.

### Genetic Burden of Rare Variants in Kidney Disease-related Genes in DKD Patients

Next, we performed gene-based burden association tests to compare the rate of rare variants between NDKD and DKD patients aggregated within each gene. As expected, the most significant enrichment of rare pathogenic or likely pathogenic variants was observed in *PKD1* among NDKD patients (*p*-value = 0.008; **Supplemental Table 6**). Similarly, although not statistically significant (*p*-value >0.05), enrichment of rare pathogenic or likely pathogenic variants was also observed in *PKD2* among NDKD patients. Conversely, an excess of rare variants in *C5orf42* was seen in DKD patients.

As variants classified as VUSs using ACMG-AMP guideline cannot be ruled out as benign and likely include pathogenic variants, we expanded our gene-based burden analyses to include rare variants with supportive evidence of pathogenicity based on computational prediction methods. As seen in our analysis of rare pathogenic or likely pathogenic variants, we observed an excess burden of rare variants in both *PKD1* (*p*-value = 0.01) and *PKD2* (*p*-value = 0.04) in NDKD patients. Rare variants in *COL4A5* (*p*-value = 0.008) were also significantly enriched in NDKD patients compared to DKD patients. Interestingly, in addition to strengthening the association seen in DKD patients with rare variants in *C5orf42* (*p*-value = 0.04), this analysis detected enrichment of rare variants in DKD patients in both *ACE* (*p*-value = 0.008) and *NEK8* (*p*-value = 0.04).

### Rare Diagnostic Variants Identified in a Subset of DKD Patients Suggest They Have NDKD

Three DKD patients (2.8%) were found to carry rare variants suggestive that the underlying cause of their kidney disease may not be a consequence of their diabetes (**Table 3**). A patient with CKD attributed to T2D, was found to have a rare diagnostic variant in *AQP2* (p.N220T); substitution mutations in this gene have been shown to cause an autosomal dominant form of nephrogenic diabetes insipidus, a condition characterized by severe polyuria (35). Similarly, a pathogenic *REN* variant (p.R49X) was identified in a second patient with T2D. Pathogenic variants in *REN* are associated with autosomal dominant tubulointerstitial kidney disease (ADTKD) (36; 37). As both of these patients also have a history of diabetic retinopathy, it is possible that they have concomitant DKD and NDKD. Lastly, 1 T2D patient was found to carry a diagnostic variant in *COL4A3* (p.R1661C). As mentioned above, this variant has been observed in patients with FSGS (31) and autosomal recessive Alport syndrome (32). In contrast to the DKD patients with *AQP2* and *REN* variants, despite more than 35 years of diabetes, this patient does not have reported evidence of retinopathy or neuropathy, further suggesting that this patient likely has NDKD.

## Discussion

The goal of this study was to investigate the utility of NGS in understanding the molecular basis of kidney disease in patients with diabetes. We successfully performed targeted NGS in a total of 206 patients with CKD, including 97 NDKD and 109 DKD patients, using a custom gene panel comprised of 345 kidney disease-related genes. Our analysis identified diagnostic variants that were consistent with the clinical diagnosis of approximately 19% of the NDKD patients included in this study. While one-third of NDKD patients were found to carry pathogenic or likely pathogenic variants, as classified by the ACMG-AMP guidelines, more than 20% of the DKD patients included in this study carried similarly classified variants (i.e., pathogenic or likely pathogenic variants) in these same kidney disease-related genes. We detected genetic variants suggestive of NDKD in approximately 3% of the diabetic patients included in this study.

Several of the genes identified in DKD patients, including *CUBN, PEX5, VDR*, and *COL4A3*, have previously been implicated in diabetic nephropathy (38–41). Coding variants in *CUBN*, a gene expressed in the apical brush border of proximal tubule cells (42), are known to be associated with albuminuria (38; 43). A recent exome-wide association study identified a rare coding variant in *CUBN* that was independent of previously identified common variants in this gene and that had greater than a 3.5-fold stronger effect in patients with diabetes relative to those without diabetes (38). Similarly, a multi-omics systems biology analysis by Saito et al. found that *PEX5*, a peroxisomal biogenesis marker, is associated with diabetic nephropathy via peroxisomal dysfunction and plays a role in the down-regulation of peroxisome-related metabolites (39). More recently, as part of the largest genome-wide association study on DKD to date, the Juvenile Diabetes Research Foundation’s Diabetic Nephropathy Collaborative Research Initiative identified 16 genome-wide significance loci associated with DKD; interestingly, 2 of the top signals from this study are in known kidney disease-related genes (*COL4A3* and *BMP7*) (40). *COL4A3* encodes a major structural component of the glomerular basement membrane and is associated with heritable nephropathies, including Alport syndrome, FSGS, and TBMN. The rare *COL4A3* variant identified in 3 patients included in our study (1 DKD patient and 2 NDKD patients) has previously been observed in a family with biopsy proven FSGS (31), suggesting that this variant may contribute to kidney disease independent of diabetes status.

In the presence of hyperglycemia, defects in cilia structure or function cause structural and functional alterations in the kidney, including podocyte effacement, interstitial inflammation, and proteinuria (44). Moreover, hyperglycemic mice with cilia dysfunction have enhanced activation of Wnt signaling and epithelial-to-mesenchymal transition, two pathways that have been implicated in DKD (45; 46). Interestingly, 25% of DKD patients positive for pathogenic or likely pathogenic variants were found to carry rare variants in known ciliopathy-associated genes, including 3 DKD patients found to harbor rare pLOF variants in *C5orf42*, a gene that is associated with rare autosomal recessive ciliopathies characterized by developmental delay, multiple congenital anomalies, and cystic kidney disease (47–49). Similarly, we identified rare, likely pathogenic variants in *XPNPEP3* in 2 DKD patients with ESRD; defects in *XPNPEP3* cause nephronophthisis-like nephropathy, a cystic kidney disease that leads to ESRD (50). Taken together, these data suggest that variants in kidney disease-related genes in the context of diabetic pathophysiology, including ciliopathy-associated genes, may play a role in the pathogenesis of kidney disease in patients with diabetes.

Previous studies examining the diagnostic utility of genetic testing in CKD have included patients across diverse nephropathy subtypes (7–12; 51). In the largest study to date, the overall diagnostic yield was found to be approximately 10%; among patients with a clinical diagnosis of diabetic nephropathy, however, the diagnostic yield was much lower, only 1.6% (9). In contrast to this study, we found that as many as 22% of DKD patients included in our study carry pathogenic or likely pathogenic in the kidney disease-related genes included on our gene panel. While the reason behind this discrepancy is unclear, it should be noted that nearly 70% of those found to have variants in our study had ESRD, suggesting, perhaps, that these variants may predispose diabetic patients to more severe kidney disease.

In line with data from previous studies, our study highlights the effectiveness of genetic testing in identifying the underlying molecular cause of disease in patients with various forms of NDKD. In addition to assigning genetic diagnoses that corroborated several NDKD patients’ initial clinical diagnoses, in a subset of patients, we were able to identify rare variants that suggested re-evaluation of their diagnoses is warranted. In particular, we identified a large homozygous deletion of *NPHP1* in 1 patient in their late 20s with stage 4 CKD who had been diagnosed with ANCA vasculitis following a renal biopsy. Interestingly, previous studies have shown that homozygous deletion of *NPHP1*, arising from homologous recombination between 45-kilobasepair repeats flanking this gene, is the most frequent mutation observed in familial juvenile nephronophthisis (MIM 256100), a rare progressive tubulointerstitial kidney disorder with autosomal recessive inheritance (28; 29). More than 85% of patients with nephronophthisis, however, are clinically diagnosed as having other nephropathies (52). Importantly, while only 25% of patients diagnosed with ANCA vasculitis develop ESRD, nearly all patients with familial juvenile nephronophthisis progress to ESRD by age 30. Since enrollment to our study, this patient has progressed to stage 5 CKD and is currently receiving renal replacement therapy. For this particular patient, genetic analysis may have aided in detecting unrecognized clinical symptoms suggested by our analysis and better informed the management of their care.

Our study has several limitations that should be acknowledged. First, our study included only 206 patients, including 109 DKD patients. Despite its modest samples size, our study is the first to comprehensively investigate the burden of rare variants in kidney disease-related genes in DKD. Not only did we find that 22% of these DKD patients carry rare variants in these genes but, for a subset of these patients, we identified a genetic diagnosis suggestive that their kidney disease was likely not a consequence of their diabetes, thereby highlighting the value of genetic screening in uncovering precise molecular diagnoses. Second, although we were able to detect both coding variants and structural rearrangements using our targeted sequencing approach, variants in non-coding regions, that potentially impact RNA expression and/or processing (e.g., those that affect transcriptional repression, exon skipping, and intron inclusion), were not evaluated in this study. While whole genome sequence (WGS) and whole transcriptome sequencing (RNA-seq) are better suited to detect such changes, both technologies are not without their own limitations. WGS, for example, presents important challenges to interpretation of the vast number of variants identified per sample. As gene expression is cell-type specific, the value of RNA-seq data, on the other hand, is limited by the availability of disease-relevant source material. Nonetheless, both WGS and RNA-seq could aid in the detection and discovery of additional disease-causing mutations in patients with kidney disease. Third, in order to increase specificity, we adhered to the ACMG-AMP standards and guidelines for variant interpretation and, as such, many of the variants identified in our study were labeled as VUSs and not correlated with clinical phenotypes. As VUSs, which are variants whose pathogenicity and function are unclear, cannot fully be ruled out as benign, this may have impacted the sensitivity of our study. Computational prediction algorithms coupled with support from functional studies could help to better understand the pathogenicity of VUSs and prioritize these for further study.

Our goal was to leverage NGS technology to better understand the genetic underpinnings of kidney disease in patients with kidney disease. Although the majority of diabetic patients included in our study were not found to have pathogenic or likely pathogenic variants in kidney disease-related genes, a nontrivial proportion, (more than 20%) were found to carry variants in kidney disease-related genes that we suspect contribute to their disease. Our findings suggest that rare variants in genes not previously linked to DKD (e.g., ciliopathy genes) contribute to DKD susceptibility. In patients with diabetes, it’s possible that hyperglycemia, in addition to variants in these genes contribute to the progression of kidney disease. Additional studies, including functional studies, are necessary to better understand the role of these variants in DKD. Genetic screening in patients with DKD can provide a molecular diagnosis of the underlying cause of kidney disease that could translate to improved precision diagnostics and aid in the prognosis and long-term management of their disease. Additionally, such information is likely to better inform patient selection for clinical trials and studies on risk factors that contribute to DKD.

## Supporting information

Supplementary Appendix Summary

## Author Contributions

JL-G and MGP conceived the idea and designed the study. JL-G, JFM, AHW, RG, SGF, MHP, and CZ researched the data. JL-G, SGF, and MGP analyzed the data with assistance from JFM and AHW. JL-G and MGP drafted the manuscript with assistance from LA and NR. All authors revised the manuscript and approved the final version. MGP is the guarantor of this work and, as such, had full access to all the data in the study and takes responsibility for the integrity of the data and the accuracy of its analysis.

## Acknowledgments

The authors acknowledge grant support from the National Kidney Foundation (MGP) and the Larry H. and Gail Miller Family Foundation Diabetes Initiative (MGP).

## References

1. U.S. Renal Data System. USRDS 2018 Annual Data Report: Atlas of End-Stage Renal Disease in the United States. Bethesda, 2018

2. Anders HJ, Huber TB, Isermann B, Schiffer M: CKD in diabetes: diabetic kidney disease versus nondiabetic kidney disease. Nat Rev Nephrol 2018;14:361–377

3. Haider DG, Peric S, Friedl A, Fuhrmann V, Wolzt M, Horl WH, Soleiman A: Kidney biopsy in patients with diabetes mellitus. Clin Nephrol 2011;76:180–185

4. Sharma SG, Bomback AS, Radhakrishnan J, Herlitz LC, Stokes MB, Markowitz GS, D’Agati VD: The modern spectrum of renal biopsy findings in patients with diabetes. Clin J Am Soc Nephrol 2013;8:1718–1724

5. Zhuo L, Ren W, Li W, Zou G, Lu J: Evaluation of renal biopsies in type 2 diabetic patients with kidney disease: a clinicopathological study of 216 cases. Int Urol Nephrol 2013;45:173–179

6. Freeman NS, Canetta PA, Bomback AS: Glomerular Diseases in Patients with Diabetes Mellitus: An Underappreciated Epidemic. Kidney360 2020;

7. Bullich G, Domingo-Gallego A, Vargas I, Ruiz P, Lorente-Grandoso L, Furlano M, Fraga G, Madrid A, Ariceta G, Borregan M, Pinero-Fernandez JA, Rodriguez-Pena L, Ballesta-Martinez MJ, Llano-Rivas I, Menica MA, Ballarin J, Torrents D, Torra R, Ars E: A kidney-disease gene panel allows a comprehensive genetic diagnosis of cystic and glomerular inherited kidney diseases. Kidney Int 2018;94:363–371

8. Connaughton DM, Kennedy C, Shril S, Mann N, Murray SL, Williams PA, Conlon E, Nakayama M, van der Ven AT, Ityel H, Kause F, Kolvenbach CM, Dai R, Vivante A, Braun DA, Schneider R, Kitzler TM, Moloney B, Moran CP, Smyth JS, Kennedy A, Benson K, Stapleton C, Denton M, Magee C, O’Seaghdha CM, Plant WD, Griffin MD, Awan A, Sweeney C, Mane SM, Lifton RP, Griffin B, Leavey S, Casserly L, de Freitas DG, Holian J, Dorman A, Doyle B, Lavin PJ, Little MA, Conlon PJ, Hildebrandt F: Monogenic causes of chronic kidney disease in adults. Kidney Int 2019;95:914–928

9. Groopman EE, Marasa M, Cameron-Christie S, Petrovski S, Aggarwal VS, Milo-Rasouly H, Li Y, Zhang J, Nestor J, Krithivasan P, Lam WY, Mitrotti A, Piva S, Kil BH, Chatterjee D, Reingold R, Bradbury D, DiVecchia M, Snyder H, Mu X, Mehl K, Balderes O, Fasel DA, Weng C, Radhakrishnan J, Canetta P, Appel GB, Bomback AS, Ahn W, Uy NS, Alam S, Cohen DJ, Crew RJ, Dube GK, Rao MK, Kamalakaran S, Copeland B, Ren Z, Bridgers J, Malone CD, Mebane CM, Dagaonkar N, Fellstrom BC, Haefliger C, Mohan S, Sanna-Cherchi S, Kiryluk K, Fleckner J, March R, Platt A, Goldstein DB, Gharavi AG: Diagnostic Utility of Exome Sequencing for Kidney Disease. N Engl J Med 2019;380:142–151

10. Lata S, Marasa M, Li Y, Fasel DA, Groopman E, Jobanputra V, Rasouly H, Mitrotti A, Westland R, Verbitsky M, Nestor J, Slater LM, D’Agati V, Zaniew M, Materna-Kiryluk A, Lugani F, Caridi G, Rampoldi L, Mattoo A, Newton CA, Rao MK, Radhakrishnan J, Ahn W, Canetta PA, Bomback AS, Appel GB, Antignac C, Markowitz GS, Garcia CK, Kiryluk K, Sanna-Cherchi S, Gharavi AG: Whole-Exome Sequencing in Adults With Chronic Kidney Disease: A Pilot Study. Ann Intern Med 2018;168:100–109

11. Mallett AJ, McCarthy HJ, Ho G, Holman K, Farnsworth E, Patel C, Fletcher JT, Mallawaarachchi A, Quinlan C, Bennetts B, Alexander SI: Massively parallel sequencing and targeted exomes in familial kidney disease can diagnose underlying genetic disorders. Kidney Int 2017;92:1493–1506

12. Wang M, Chun J, Genovese G, Knob AU, Benjamin A, Wilkins MS, Friedman DJ, Appel GB, Lifton RP, Mane S, Pollak MR: Contributions of Rare Gene Variants to Familial and Sporadic FSGS. J Am Soc Nephrol 2019;30:1625–1640

13. Levey AS, Stevens LA, Schmid CH, Zhang YL, Castro AF, 3rd, Feldman HI, Kusek JW, Eggers P, Van Lente F, Greene T, Coresh J, Ckd EPI: A new equation to estimate glomerular filtration rate. Ann Intern Med 2009;150:604–612

14. Rohland N, Reich D: Cost-effective, high-throughput DNA sequencing libraries for multiplexed target capture. Genome Res 2012;22:939–946

15. Manichaikul A, Mychaleckyj JC, Rich SS, Daly K, Sale M, Chen WM: Robust relationship inference in genome-wide association studies. Bioinformatics 2010;26:2867–2873

16. Wang K, Li M, Hakonarson H: ANNOVAR: functional annotation of genetic variants from high-throughput sequencing data. Nucleic Acids Res 2010;38:e164

17. Karczewski KJ, Francioli LC, Tiao G, Cummings BB, Alföldi J, Wang Q, Collins RL, Laricchia KM, Ganna A, Birnbaum DP, Gauthier LD, Brand H, Solomonson M, Watts NA, Rhodes D, Singer-Berk M, England EM, Seaby EG, Kosmicki JA, Walters RK, Tashman K, Farjoun Y, Banks E, Poterba T, Wang A, Seed C, Whiffin N, Chong JX, Samocha KE, Pierce-Hoffman E, Zappala Z, O’Donnell-Luria AH, Minikel EV, Weisburd B, Lek M, Ware JS, Vittal C, Armean IM, Bergelson L, Cibulskis K, Connolly KM, Covarrubias M, Donnelly S, Ferriera S, Gabriel S, Gentry J, Gupta N, Jeandet T, Kaplan D, Llanwarne C, Munshi R, Novod S, Petrillo N, Roazen D, Ruano-Rubio V, Saltzman A, Schleicher M, Soto J, Tibbetts K, Tolonen C, Wade G, Talkowski ME,, Neale BM, Daly MJ, MacArthur DG: Variation across 141,456 human exomes and genomes reveals the spectrum of loss-of-function intolerance across human proteincoding genes. bioRxiv 2019;

18. Li Q, Wang K: InterVar: Clinical Interpretation of Genetic Variants by the 2015 ACMG-AMP Guidelines. Am J Hum Genet 2017;100:267–280

19. Richards S, Aziz N, Bale S, Bick D, Das S, Gastier-Foster J, Grody WW, Hegde M, Lyon E, Spector E, Voelkerding K, Rehm HL, Committee ALQA: Standards and guidelines for the interpretation of sequence variants: a joint consensus recommendation of the American College of Medical Genetics and Genomics and the Association for Molecular Pathology. Genet Med 2015;17:405–424

20. Sim NL, Kumar P, Hu J, Henikoff S, Schneider G, Ng PC: SIFT web server: predicting effects of amino acid substitutions on proteins. Nucleic Acids Res 2012;40:W452–457

21. Adzhubei IA, Schmidt S, Peshkin L, Ramensky VE, Gerasimova A, Bork P, Kondrashov AS, Sunyaev SR: A method and server for predicting damaging missense mutations. Nat Methods 2010;7:248–249

22. Schwarz JM, Rodelsperger C, Schuelke M, Seelow D: MutationTaster evaluates disease-causing potential of sequence alterations. Nat Methods 2010;7:575–576

23. Jagadeesh KA, Wenger AM, Berger MJ, Guturu H, Stenson PD, Cooper DN, Bernstein JA, Bejerano G: M-CAP eliminates a majority of variants of uncertain significance in clinical exomes at high sensitivity. Nat Genet 2016;48:1581–1586

24. Chun S, Fay JC: Identification of deleterious mutations within three human genomes. Genome Res 2009;19:1553–1561

25. Chiang T, Liu X, Wu TJ, Hu J, Sedlazeck FJ, White S, Schaid D, Andrade M, Jarvik GP, Crosslin D, Stanaway I, Carrell DS, Connolly JJ, Hakonarson H, Groopman EE, Gharavi AG, Fedotov A, Bi W, Leduc MS, Murdock DR, Jiang Y, Meng L, Eng CM, Wen S, Yang Y, Muzny DM, Boerwinkle E, Salerno W, Venner E, Gibbs RA: Atlas-CNV: a validated approach to call single-exon CNVs in the eMERGESeq gene panel. Genet Med 2019;21:2135–2144

26. Robinson JT, Thorvaldsdottir H, Winckler W, Guttman M, Lander ES, Getz G, Mesirov JP: Integrative genomics viewer. Nat Biotechnol 2011;29:24–26

27. Lee S, Emond MJ, Bamshad MJ, Barnes KC, Rieder MJ, Nickerson DA, Team NGESP-ELP, Christiani DC, Wurfel MM, Lin X: Optimal unified approach for rare-variant association testing with application to small-sample case-control whole-exome sequencing studies. Am J Hum Genet 2012;91:224–237

28. Konrad M, Saunier S, Heidet L, Silbermann F, Benessy F, Calado J, Le Paslier D, Broyer M, Gubler MC, Antignac C: Large homozygous deletions of the 2q13 region are a major cause of juvenile nephronophthisis. Hum Mol Genet 1996;5:367–371

29. Saunier S, Calado J, Benessy F, Silbermann F, Heilig R, Weissenbach J, Antignac C: Characterization of the NPHP1 locus: mutational mechanism involved in deletions in familial juvenile nephronophthisis. Am J Hum Genet 2000;66:778–789

30. Holmes LV, Strain L, Staniforth SJ, Moore I, Marchbank K, Kavanagh D, Goodship JA, Cordell HJ, Goodship TH: Determining the population frequency of the CFHR3/CFHR1 deletion at 1q32. PLoS One 2013;8:e60352

31. Malone AF, Phelan PJ, Hall G, Cetincelik U, Homstad A, Alonso AS, Jiang R, Lindsey TB, Wu G, Sparks MA, Smith SR, Webb NJ, Kalra PA, Adeyemo AA, Shaw AS, Conlon PJ, Jennette JC, Howell DN, Winn MP, Gbadegesin RA: Rare hereditary COL4A3/COL4A4 variants may be mistaken for familial focal segmental glomerulosclerosis. Kidney Int 2014;86:1253–1259

32. Heidet L, Arrondel C, Forestier L, Cohen-Solal L, Mollet G, Gutierrez B, Stavrou C, Gubler MC, Antignac C: Structure of the human type IV collagen gene COL4A3 and mutations in autosomal Alport syndrome. J Am Soc Nephrol 2001;12:97–106

33. Terryn W, Vanholder R, Hemelsoet D, Leroy BP, Van Biesen W, De Schoenmakere G, Wuyts B, Claes K, De Backer J, De Paepe G, Fogo A, Praet M, Poppe B: Questioning the Pathogenic Role of the GLA p.Ala143Thr “Mutation” in Fabry Disease: Implications for Screening Studies and ERT. JIMD Rep 2013;8:101–108

34. Takahara M, Katoh Y, Nakamura K, Hirano T, Sugawa M, Tsurumi Y, Nakayama K: Ciliopathy-associated mutations of IFT122 impair ciliary protein trafficking but not ciliogenesis. Hum Mol Genet 2018;27:516–528

35. Loonen AJ, Knoers NV, van Os CH, Deen PM: Aquaporin 2 mutations in nephrogenic diabetes insipidus. Semin Nephrol 2008;28:252–265

36. Schaeffer C, Izzi C, Vettori A, Pasqualetto E, Cittaro D, Lazarevic D, Caridi G, Gnutti B, Mazza C, Jovine L, Scolari F, Rampoldi L: Autosomal Dominant Tubulointerstitial Kidney Disease with Adult Onset due to a Novel Renin Mutation Mapping in the Mature Protein. Sci Rep 2019;9:11601

37. Zivna M, Hulkova H, Matignon M, Hodanova K, Vylet’al P, Kalbacova M, Baresova V, Sikora J, Blazkova H, Zivny J, Ivanek R, Stranecky V, Sovova J, Claes K, Lerut E, Fryns JP, Hart PS, Hart TC, Adams JN, Pawtowski A, Clemessy M, Gasc JM, Gubler MC, Antignac C, Elleder M, Kapp K, Grimbert P, Bleyer AJ, Kmoch S: Dominant renin gene mutations associated with early-onset hyperuricemia, anemia, and chronic kidney failure. Am J Hum Genet 2009;85:204–213

38. Ahluwalia TS, Schulz CA, Waage J, Skaaby T, Sandholm N, van Zuydam N, Charmet R, Bork-Jensen J, Almgren P, Thuesen BH, Bedin M, Brandslund I, Christensen CK, Linneberg A, Ahlqvist E, Groop PH, Hadjadj S, Tregouet DA, Jorgensen ME, Grarup N, Pedersen O, Simons M, Groop L, Orho-Melander M, McCarthy MI, Melander O, Rossing P, Kilpelainen TO, Hansen T: A novel rare CUBN variant and three additional genes identified in Europeans with and without diabetes: results from an exome-wide association study of albuminuria. Diabetologia 2019;62:292–305

39. Saito R, Rocanin-Arjo A, You YH, Darshi M, Van Espen B, Miyamoto S, Pham J, Pu M, Romoli S, Natarajan L, Ju W, Kretzler M, Nelson R, Ono K, Thomasova D, Mulay SR, Ideker T, D’Agati V, Beyret E, Belmonte JCI, Anders HJ, Sharma K: Systems biology analysis reveals role of MDM2 in diabetic nephropathy. JCI Insight 2016;1:e87877

40. Salem RM, Todd JN, Sandholm N, Cole JB, Chen WM, Andrews D, Pezzolesi MG, McKeigue PM, Hiraki LT, Qiu C, Nair V, Di Liao C, Cao JJ, Valo E, Onengut-Gumuscu S, Smiles AM, McGurnaghan SJ, Haukka JK, Harjutsalo V, Brennan EP, van Zuydam N, Ahlqvist E, Doyle R, Ahluwalia TS, Lajer M, Hughes MF, Park J, Skupien J, Spiliopoulou A, Liu A, Menon R, Boustany-Kari CM, Kang HM, Nelson RG, Klein R, Klein BE, Lee KE, Gao X, Mauer M, Maestroni S, Caramori ML, de Boer IH, Miller RG, Guo J, Boright AP, Tregouet D, Gyorgy B, Snell-Bergeon JK, Maahs DM, Bull SB, Canty AJ, Palmer CNA, Stechemesser L, Paulweber B, Weitgasser R, Sokolovska J, Rovite V, Pirags V, Prakapiene E, Radzeviciene L, Verkauskiene R, Panduru NM, Groop LC, McCarthy MI, Gu HF, Mollsten A, Falhammar H, Brismar K, Martin F, Rossing P, Costacou T, Zerbini G, Marre M, Hadjadj S, McKnight AJ, Forsblom C, McKay G, Godson C, Maxwell AP, Kretzler M, Susztak K, Colhoun HM, Krolewski A, Paterson AD, Groop PH, Rich SS, Hirschhorn JN, Florez JC, Summit Consortium DERGGC: Genome-Wide Association Study of Diabetic Kidney Disease Highlights Biology Involved in Glomerular Basement Membrane Collagen. J Am Soc Nephrol 2019;30:2000–2016

41. Sanchez-Nino MD, Bozic M, Cordoba-Lanus E, Valcheva P, Gracia O, Ibarz M, Fernandez E, Navarro-Gonzalez JF, Ortiz A, Valdivielso JM: Beyond proteinuria: VDR activation reduces renal inflammation in experimental diabetic nephropathy. Am J Physiol Renal Physiol 2012;302:F647–657

42. Amsellem S, Gburek J, Hamard G, Nielsen R, Willnow TE, Devuyst O, Nexo E, Verroust PJ, Christensen EI, Kozyraki R: Cubilin is essential for albumin reabsorption in the renal proximal tubule. J Am Soc Nephrol 2010;21:1859–1867

43. Boger CA, Chen MH, Tin A, Olden M, Kottgen A, de Boer IH, Fuchsberger C, O’Seaghdha CM, Pattaro C, Teumer A, Liu CT, Glazer NL, Li M, O’Connell JR, Tanaka T, Peralta CA, Kutalik Z, Luan J, Zhao JH, Hwang SJ, Akylbekova E, Kramer H, van der Harst P, Smith AV, Lohman K, de Andrade M, Hayward C, Kollerits B, Tonjes A, Aspelund T, Ingelsson E, Eiriksdottir G, Launer LJ, Harris TB, Shuldiner AR, Mitchell BD, Arking DE, Franceschini N, Boerwinkle E, Egan J, Hernandez D, Reilly M, Townsend RR, Lumley T, Siscovick DS, Psaty BM, Kestenbaum B, Haritunians T, Bergmann S, Vollenweider P, Waeber G, Mooser V, Waterworth D, Johnson AD, Florez JC, Meigs JB, Lu X, Turner ST, Atkinson EJ, Leak TS, Aasarod K, Skorpen F, Syvanen AC, Illig T, Baumert J, Koenig W, Kramer BK, Devuyst O, Mychaleckyj JC, Minelli C, Bakker SJ, Kedenko L, Paulweber B, Coassin S, Endlich K, Kroemer HK, Biffar R, Stracke S, Volzke H, Stumvoll M, Magi R, Campbell H, Vitart V, Hastie ND, Gudnason V, Kardia SL, Liu Y, Polasek O, Curhan G, Kronenberg F, Prokopenko I, Rudan I, Arnlov J, Hallan S, Navis G, Consortium CK, Parsa A, Ferrucci L, Coresh J, Shlipak MG, Bull SB, Paterson NJ, Wichmann HE, Wareham NJ, Loos RJ, Rotter JI, Pramstaller PP, Cupples LA, Beckmann JS, Yang Q, Heid IM, Rettig R, Dreisbach AW, Bochud M, Fox CS, Kao WH: CUBN is a gene locus for albuminuria. J Am Soc Nephrol 2011;22:555–570

44. Sas KM, Yin H, Fitzgibbon WR, Baicu CF, Zile MR, Steele SL, Amria M, Saigusa T, Funk J, Bunni MA, Siegal GP, Siroky BJ, Bissler JJ, Bell PD: Hyperglycemia in the absence of cilia accelerates cystogenesis and induces renal damage. Am J Physiol Renal Physiol 2015;309:F79–87

45. Hills CE, Squires PE: The role of TGF-beta and epithelial-to mesenchymal transition in diabetic nephropathy. Cytokine Growth Factor Rev 2011;22:131–139

46. Zhou T, He X, Cheng R, Zhang B, Zhang RR, Chen Y, Takahashi Y, Murray AR, Lee K, Gao G, Ma JX: Implication of dysregulation of the canonical wingless-type MMTV integration site (WNT) pathway in diabetic nephropathy. Diabetologia 2012;55:255–266

47. Fleming LR, Doherty DA, Parisi MA, Glass IA, Bryant J, Fischer R, Turkbey B, Choyke P, Daryanani K, Vemulapalli M, Mullikin JC, Malicdan MC, Vilboux T, Sayer JA, Gahl WA, Gunay-Aygun M: Prospective Evaluation of Kidney Disease in Joubert Syndrome. Clin J Am Soc Nephrol 2017;12:1962–1973

48. Romani M, Mancini F, Micalizzi A, Poretti A, Miccinilli E, Accorsi P, Avola E, Bertini E, Borgatti R, Romaniello R, Ceylaner S, Coppola G, D’Arrigo S, Giordano L, Janecke AR, Lituania M, Ludwig K, Martorell L, Mazza T, Odent S, Pinelli L, Poo P, Santucci M, Signorini S, Simonati A, Spiegel R, Stanzial F, Steinlin M, Tabarki B, Wolf NI, Zibordi F, Boltshauser E, Valente EM: Oral-facial-digital syndrome type VI: is C5orf42 really the major gene? Hum Genet 2015;134:123–126

49. Wentzensen IM, Johnston JJ, Keppler-Noreuil K, Acrich K, David K, Johnson KD, Graham JM, Jr., Sapp JC, Biesecker LG: Exome sequencing identifies novel mutations in C5orf42 in patients with Joubert syndrome with oral-facial-digital anomalies. Hum Genome Var 2015;2:15045

50. O’Toole JF, Liu Y, Davis EE, Westlake CJ, Attanasio M, Otto EA, Seelow D, Nurnberg G, Becker C, Nuutinen M, Karppa M, Ignatius J, Uusimaa J, Pakanen S, Jaakkola E, van den Heuvel LP, Fehrenbach H, Wiggins R, Goyal M, Zhou W, Wolf MT, Wise E, Helou J, Allen SJ, Murga-Zamalloa CA, Ashraf S, Chaki M, Heeringa S, Chernin G, Hoskins BE, Chaib H, Gleeson J, Kusakabe T, Suzuki T, Isaac RE, Quarmby LM, Tennant B, Fujioka H, Tuominen H, Hassinen I, Lohi H, van Houten JL, Rotig A, Sayer JA, Rolinski B, Freisinger P, Madhavan SM, Herzer M, Madignier F, Prokisch H, Nurnberg P, Jackson PK, Khanna H, Katsanis N, Hildebrandt F: Individuals with mutations in XPNPEP3, which encodes a mitochondrial protein, develop a nephronophthisis-like nephropathy. J Clin Invest 2010;120:791–802

51. Renkema KY, Stokman MF, Giles RH, Knoers NV: Next-generation sequencing for research and diagnostics in kidney disease. Nat Rev Nephrol 2014;10:433–444

52. Snoek R, van Setten J, Keating BJ, Israni AK, Jacobson PA, Oetting WS, Matas AJ, Mannon RB, Zhang Z, Zhang W, Hao K, Murphy B, Reindl-Schwaighofer R, Heinzl A, Oberbauer R, Viklicky O, Conlon PJ, Stapleton CP, Bakker SJL, Snieder H, Peters EDJ, van der Zwaag B, Knoers N, De Borst MH, van Eerde AM: NPHP1 (Nephrocystin-1) Gene Deletions Cause Adult-Onset ESRD. J Am Soc Nephrol 2018;29:1772–1779

