## Supplementary Appendix Summary for "Targeted Next-Generation Sequencing Identifies Pathogenic Variants in Kidney Disease-Related Genes in Patients with Diabetic Kidney Disease"

This appendix has been provided by the authors to give readers additional information about their work.

[**Table S4. Rare (MAF <0.1%), Pathogenic/Likely Pathogenic, and VUS.**	7](#_Toc35728746)

[**Table S5. Rare (MAF <0.1%), Pathogenic/Likely Pathogenic, and VUS Biallelic variants per Patients.**	8](#_Toc35728747)

**Table S1. Utah Kidney Gene Panel.**

345 listed genes, including protein accession number, isoforms number, OMIM accession number and inheritance mode.

See **Supplementary Appendix.xlsx** on Diabetes.diabetesjournals.org: sheet “**TableS1**”

OMIM: Online Mendelian Inheritance of Man.

| Variants class | All Variants | MAF <0.1%* | MAF < 0.1% ACMG-AMP Classification | | | | |
| --- | --- | --- | --- | --- | --- | --- | --- |
|  |  |  | Pathogenic | Likely Pathogenic | VUS | Likely Benign | Benign |
| Nonsynonymous | 2,673 | 1,156 | 0 | 19 | 968 | 157 | 12 |
| Stop Gain | 28 | 24 | 12 | 3 | 9 | 0 | 0 |
| Stop Loss | 2 | 1 | 0 | 0 | 1 | 0 | 0 |
| Splicing | 23 | 13 | 9 | 0 | 4 | 0 | 0 |
| Frameshift Deletion | 25 | 20 | 2 | 12 | 6 | 0 | 0 |
| Frameshift Insertion | 17 | 9 | 1 | 5 | 3 | 0 | 0 |
| In-frame Deletion | 44 | 25 | 0 | 0 | 23 | 2 | 0 |
| In-frame insertion | 24 | 11 | 0 | 0 | 10 | 0 | 1 |
| Total | 2,836 | 1,259 | 24 | 39 | 1,024 | 159 | 13 |

**Table S2: Summary of Identified Functional Variants**

MAF, minor allele frequency (*based on gnomAD MAF); ACMG-AMP, American College of Medical Genetics and Genomics and the Association for Molecular Pathology; VUS, variant of unknown significance.

**Table S3. Variants Identified in 345 Kidney-disease Related Genes in NDKD and DKD Patients.**

Variants Identified in 345 Kidney-disease Related Genes in NDKD and DKD Patients. A total of 2,836 functional variants (nonsynonymous, stop gain, stop loss, splicing, frameshift deletion/insertion, and in-frame deletion/insertion were identified in 202 patients.

\# Damaging (Count): the number of algorithms predicting the listed variant as damaging by the SIFT, LRT, MutationTaster, PolyPhen2, and MCAP algorithms.

See **Supplementary Appendix.xlsx** on Diabetes.diabetesjournals.org: sheet “**TableS3**”

**Table S4. Rare (MAF <0.1%), Pathogenic/Likely Pathogenic, and VUS.**

A total of 24 pathogenic variants, 39 likely pathogenic variants, and 1,024 VUSs were identified in 202 patients.

\# Damaging (Count): the number of algorithms predicting the listed variant as damaging by the SIFT, LRT, MutationTaster, PolyPhen2, and MCAP algorithms.

See **Supplementary Appendix.xlsx** on Diabetes.diabetesjournals.org: sheet “**TableS4**”

**Table S5. Rare (MAF <0.1%), Pathogenic/Likely Pathogenic, and VUS Biallelic variants per Patients.**

A total of 77 variants, 3 Pathogenic, 4 likely pathogenic variants, and 70 VUSs were identified in 30 patients.

\# Damaging (Count): the number of algorithms predicting the listed variant as damaging by the SIFT, LRT, MutationTaster, PolyPhen2, and MCAP algorithms.

A total of 24 pathogenic variants, 39 likely pathogenic variants, and 1,024 VUSs were identified in 202 patients. # Damaging (Count): the number of algorithms predicting the listed variant as damaging by the SIFT, LRT, MutationTaster, PolyPhen2, and MCAP algorithms.

See **Supplementary Appendix.xlsx** on Diabetes.diabetesjournals.org: sheet “**TableS5**”

**Table S6: Gene-Based Burden Association Tests (SKAT) of Rare Variants in NDKD and DKD Patients**

| Rare Pathogenic and Likely Pathogenic Variants: | | |
| --- | --- | --- |
| Gene | Number of Rare Variants | *P*-value |
| *PKD1* | 6 | 0.008 |
| *C5orf42* | 3 | 0.09 |
| *PKD2* | 3 | 0.09 |
| Rare Pathogenic, Likely Pathogenic, Variants with Supportive Evidence of Pathogenicity: | | |
| Gene | Number of Rare Variants | *P*-value |
| *ACE* | 5 | 0.008 |
| *COL4A5* | 6 | 0.008 |
| *PKD1* | 15 | 0.01 |
| *C5orf42* | 5 | 0.04 |
| *PKD2* | 4 | 0.04 |
| *NEK8* | 4 | 0.04 |

SKAT: SNP-set (Sequence) Kernel Association Test.

**Figure S1: *NPHP1* Gene Homozygous Deletion in One Patient in NDKD Cohort.**

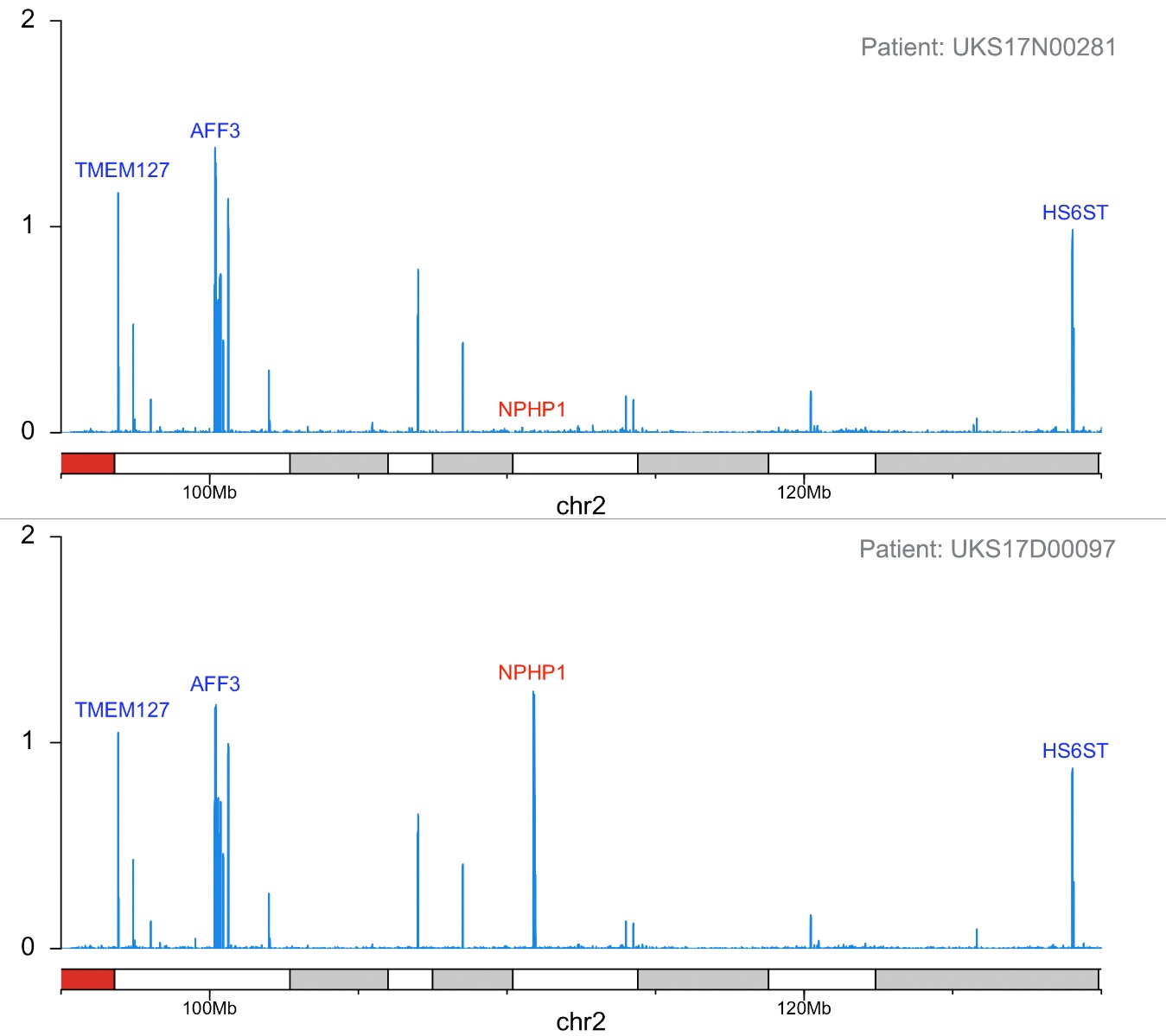

Top: CNV analysis from a NDKD patient with a deletion of *NPHP1* gene.

Bottom: CNV analysis from a NDKD patient without a deletion of *NPHP1* gene.

NDKD: Non-Diabetic Kidney Diseases

CNV: Copy number variant
